# A spatially-explicit model of stabilizing selection for improving phylogenetic inference

**DOI:** 10.1101/2020.05.12.091744

**Authors:** Jeremy M. Beaulieu, Brian C. O’Meara, Michael A. Gilchrist

**Affiliations:** Department of Biological Sciences, University of Arkansas, Fayetteville, Arkansas, 72701 USA; Department of Ecology and Evolutionary Biology, University of Tennessee, Knoxville, Tennessee, 37996-1610 USA

**Keywords:** Ultraconserved elements, Wright-Fisher, stabilizing selection, mutation

## Abstract

Ultraconserved elements (UCEs) are stretches of hundreds of nucleotides with highly conserved cores flanked by variable regions. Although the selective forces responsible for the preservation of UCEs are unknown, they are nonetheless believed to contain phylogenetically meaningful information from deep to shallow divergence events. Phylogenetic applications of UCEs assume the same degree of rate heterogeneity applies across the entire locus, including variable flanking regions. We present a Wright-Fisher model of selection on nucleotides (SelON) which includes the effects of mutation, drift, and spatially varying, stabilizing selection for an optimal nucleotide sequence. The SelON model assumes the strength of stabilizing selection follows a position dependent Gaussian function whose exact shape can vary between UCEs. We evaluate SelON by comparing its performance to a simpler and spatially invariant GTR+*Γ* model using an empirical dataset of 400 vertebrate UCEs used to determine the phylogenetic position of turtles. We observe much improvement in model fit of SelON over the GTR+*Γ* model, and support for turtles as sister to lepidosaurs. Overall, the UCE specific parameters SelON estimates provide a compact way of quantifying the strength and variation in selection within and across UCEs. SelON can also be extended to include more realistic mapping functions between sequence and stabilizing selection as well as allow for greater levels of rate heterogeneity. By more explicitly modeling the nature of selection on UCEs, SelON and similar approaches can be used to better understand the biological mechanisms responsible for their preservation across highly divergent taxa and long evolutionary time scales.

## Introduction

High-throughput DNA sequencing has transformed phylogenetics from individual targeting of a subset of genes to genome-scale approaches for the simultaneous sequence capture of entire genomes. However, there are important technical challenges when analyzing genomes across a set of species. This has led to the development of reduced representation approaches such as restriction-site associated DNA markers sequencing (e.g., Miller et al. 2007), or targeted capture of entire organellar genomes (e.g., Cronn et al. 2008), select protein-coding genes (e.g., exome capture; Hodges et al. 2007), and ultraconserved genomic elements (i.e., UCEs; Bejerano et al. 2004) that specifically target only the informative portions of the genome. The end result is still an extraordinary wealth of data, which holds great promise for resolving the tree of life.

With these large phylogenomic datasets also comes opportunities for gaining new and important insights about the evolutionary processes occurring within the genome. For example, as their name implies, UCEs are particularly unique in that they are highly conserved across divergent taxa. However, because UCEs are so strongly conserved over a wide range of both time and taxa, and because they may, in some cases, serve critical regulatory functions within the genome, there is evidence that they are subject to strong stabilizing selection (e.g., Berejano et al. 2004; Woolfe et al. 2005; Katzman et al. 2007). The increasingly variable regions flanking each UCE suggests that the sensitivity of the element to selection changes based on nucleotide position (Faircloth et al. 2012; Van Dam et al. 2017). While there have been advances such as automated pipelines for identifying partitioning schemes (Tagliacollo and Lanfear 2018), our goal is to contribute to developing a more mechanistic understanding of UCE and their evolution, in addition to their utility in phylogenetic inference, by explicitly modeling the spatial variation in selection hypothesized to be responsible for UCEs.

Here, we model nucleotide substitution across a UCE as a Wright-Fisher process, which includes the processes of stabilizing selection, mutation, and drift, to link fitness to a binary distance function between an observed nucleotide and the “optimal” base for a site. With our modeling approach, unique continuous functions scale each UCE, governing how the strength of selection changes across nucleotide position, which provides a more realistic, spatially explicit model that exists outside the classic substitution-based framework. We implement and test our model using simulations, and we also apply it to a well-cited empirical data set (Crawford et al. 2012) which addressed the phylogenetic position of turtles relative to archosaurs (i.e., birds + crocodiles) and lepidosaurs (i.e., squamates + tuataras). Unlike Crawford et al (2012), who used standard substitution-based models, with our model we find that placing the turtles sister to lepidosaurs is substantially better supported than placing turtles sister to archosaurs. Similar to Crawford et al. (2012), we find that the estimated length of the branches that distinguishes the two rival hypotheses to be extremely short. We further explore the information content contained both among and within each UCE and test whether particular attributes of the UCEs themselves or whether how the rate distribution is structured across sites, correlates with favoring one topology over the other. On the whole, we show how the UCE specific parameters estimated by our model provides a novel way of quantifying the strength and variation in selection operating within and across UCE loci.

### New Modeling Approach

We begin by constructing the global mutation matrix, **M**, whose elements define the mutation rate, *μ_ij_*, from nucleotides *i* and *j* across a set of UCEs. For our purposes, we rely on the general unrestricted model (referred to hereafter as UNREST; see Yang 1994), because it does not impose any constraints on the instantaneous rates of change. However, more constrained nucleotide models could be used instead, ranging from Jukes-Cantor (JC69; Jukes and Cantor 1969) to the Hasegawa-Kishino-Yano (HKY85, Hasegawa et al. 1985) to the Generalized time-reversible (GTR, Taveré 1986) substitution models. Since we are defining the relative rates of change, we arbitrarily set the G→T mutation rate to 1, with the remaining 11 mutation rate parameters freely estimated. As with any standard nucleotide model, the diagonals *μ_ii_* are set to −**∑**_*j*≠*i*_ *μ_i,j_*. We assume that the mutation matrix, **M**, is shared across all UCEs.

Following Sella and Hirsh (2005), we allow the processes of selection and drift to come into the model through the construction of a second matrix, **U**, whose entries, *u_ij_* describe the fixation probability of a nucleotide *j* introduced via mutation into a resident population of nucleotide *i*. First, we let *x_k_* represent the nucleotide position within the element, and let *f*(*x*) be a continuous function that describes the strength of selection on the element to match a particular optimal target nucleotide sequence 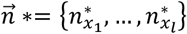, where *l* represents the length of the element. For our purposes, we assume that 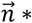 is fixed; that is, the optimal target sequence does not change along the tree. As we discuss later, this assumption could be relaxed in future models, but is, to a rough approximation, consistent with the definitional behavior of UCEs. We let 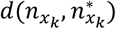 be a function that measures the functional distance between the nucleotide at position *x_k_, n_x_k__*, for a given observed nucleotide and the target nucleotide at that same position, 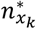. In other contexts, we could treat this as a continuous distance (e.g., Goldman and Yang 1994; Beaulieu et al. 2019), but for now, we treat the functional distance as a binary function such that,

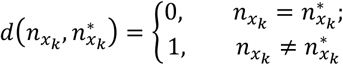

As mentioned above, the shape of the potential rate distribution (i.e., site similarity by nucleotide position; also see Faircloth et al. 2012; Van Dam et al. 2017) for a given UCE implies that the strength of selection changes with position. Thus, we define a continuous function, *f*(*x*), to describe how this sensitivity changes with nucleotide position. When *f*(*x*) is large, observed sequences that differ from *n*\* at that position are assumed under strong stabilizing selection. By contrast, when *f*(*x*) is small, then sequences that differ from *n*\* at that position are under very weak stabilizing selection. Since the form of *f*(*x*) is unknown *a priori*, we model site-specific sensitivity to selection according to a Gaussian function,

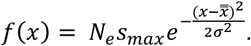

Specifically, we estimate *N_e_s_max_*, which defines the maxima of the curve, 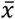, which defines the position of the center of the distribution, and *σ* which sets the width of the distribution (Fig. 1B). Because we estimate 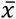, the Gaussian function need not always assume that the conserved portion is centered, and thus can and will produce a variety of shapes (see Fig. 2).

**Figure 1.**
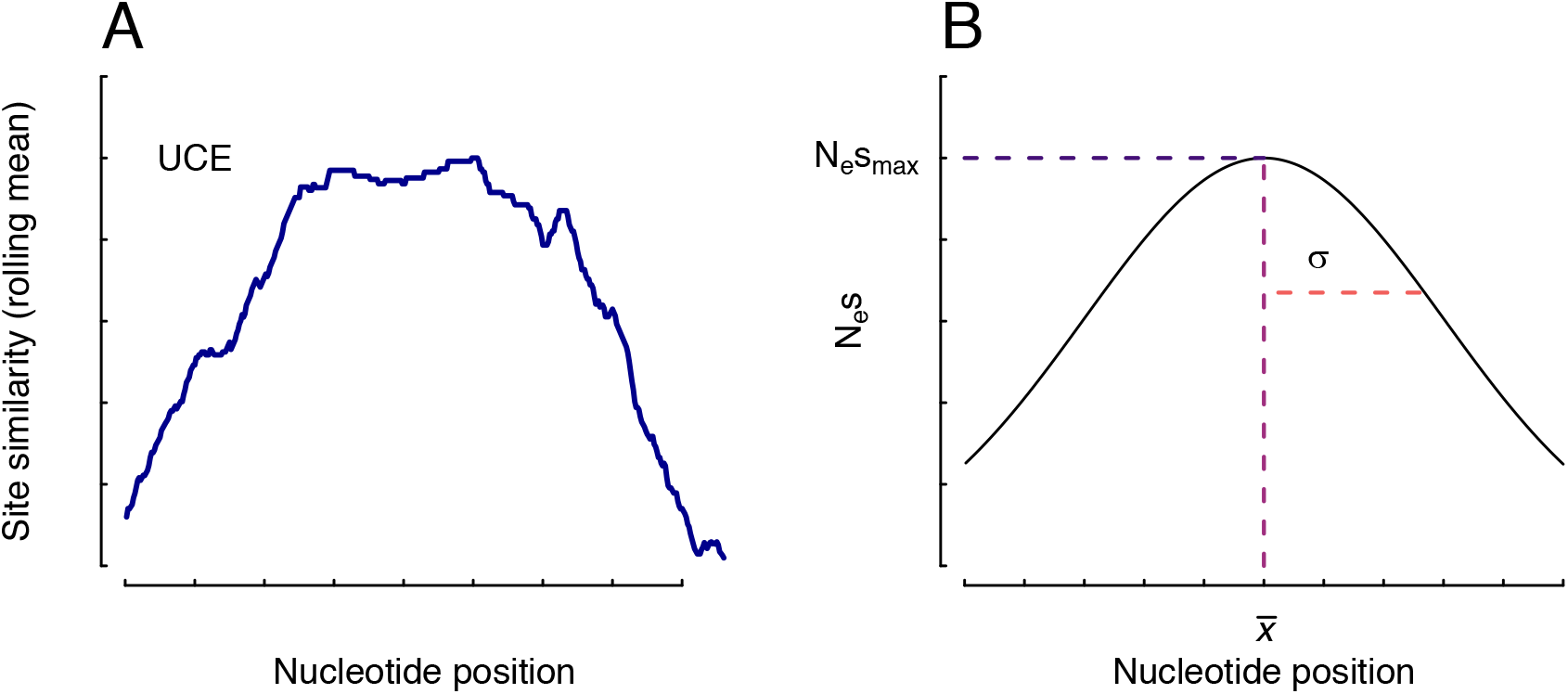
(A) An empirical example of the rate distribution across an ultra-conserved element (UCE), using site-similarity (i.e., the average distance between all taxon pairs at a site) as a proxy for rate. (B) A conceptual diagram of the generic Gaussian function used to model sensitivity to selection. The parameter *N_e_s_max_*, defines the maxima of the curve, 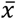, defines the site position of the center of the distribution, and *σ* sets the width of the distribution. The Gaussian distribution best approximates the overall shape of the rate variation where the strongly conserved portion of the ultraconserved element (UCE) are under strong selection, flanked on either side by highly variable stretches that are less sensitive to selection.

Assuming that the contributions to fitness are independent between nucleotide positions, we define the fitness of an observed sequence given an optimal sequence 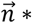 as,

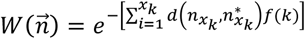

Assuming weak mutation (i.e., *N_e_* << 1), the fixation probability of a newly introduced allele depends critically on *N_e_* and the genotype fitness ratios. In our model, the genotype fitness ratios can be written as,

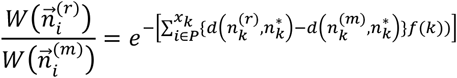

where 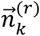 is the resident allele and 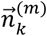 is the newly introduced mutant allele and *P* is the set of nucleotide positions where the two genotypes differ. As a result, we can now fill out the entries in **U** that define the probability that new mutant *j* introduced via mutation into a resident population, *i*, with effective population size *N_e_*, will go to fixation,

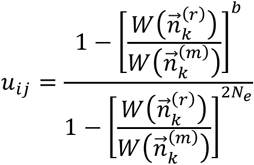

where *b*=1 for a diploid population, and *b*=2 for a haploid population.

Finally, we bring the elements of **M** and **U** together to form a single, site-specific substitution rate matrix, **Q***_x_k__*. Specifically, the elements in **Q***_x_k__* at position *x_k_* define the substitution rate from nucleotide *i* to *j* as,

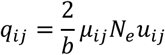

We then scale **Q***_x_k__* by a diagonal matrix **Π***_k_k__*, whose entries *π_ii_*, correspond to the equilibrium frequencies of each base at position *x_k_*. The equilibrium frequencies are determined by solving **Π***_x_k__***Q***_x_k__*=0. The final substitution matrix **Q***_x_k__* is then multiplied by the reciprocal of a scale factor, *C* = −∑*_i_π_ii_q_ii_*, to ensure that at equilibrium, one unit of branch length represents one expected substitution per site (see Fig S1). The end result is that each site has its own unique **Q***_x_k__* and equilibrium frequency, which assumes a constant mutation rate and incorporates sensitivity to selection for the target nucleotide at that given position. Because we model the evolution of each site in an independent manner, we can use the standard likelihood formula, *L_x_k__* = *P*(*D*^(*x_k_*)^|**Q***_x_k__*,*T*), for observing a site pattern, **Q***^(x_k_)^*, at position *x_k_* given the sitespecific **Q***_x_k__*, and a fixed topology and a set of branch lengths (denoted by *T*). The overall likelihood of the entire element is simply the product of the site likelihoods across *l* nucleotide positions. We do note, however, that while evolution among sites is technically independent, the shape defined by *f*(*x*) necessarily links sensitivities to selection among neighboring sites, forcing a kind of autocorrelation in rates among sites (see Fig. 1B). Overall, the log of the likelihood is maximized by estimating the genomescale mutation parameters defined by **M**, the continuous shape parameters (i.e., 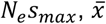, and *σ*), and the optimal nucleotide, 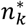, for each position of the element.

## Results

Simulations were performed to assess the difficulties in estimating the model parameters contained within our selection model (we will refer to our model hereafter simply as SelON, which stands for selection on nucleotides). In addition, these simulations were designed to determine the behavior of standard models of nucleotide substitution, like the general time-reversible (GTR) model with Gammadistributed (+*Γ*) rate variation, when sequences are generated under SelON. With regards to the parameter estimation under SelON, the model can recover the known values from the generating model quite well. Among the set of simulated UCEs the individual shapes of the strength of selection varied considerably (Fig. 2, Fig. S2-S3), and SelON properly estimated the centering, width, and magnitude of these shapes with generally very little uncertainty among the simulation replicates (Fig. 2, Fig. S2-S3). However, even though the overall shapes closely followed those contained within the generating model, there was a slight upward bias in estimates of the individual width parameter, *σ,* which was consistent among the two scenarios we tested. This upward bias in our MLE of σ is consistent with a general bias in estimating the mean and variance of a normal distribution. The parameters that define the global mutation matrix, **M**, were estimated quite well, although this is somewhat unsurprising given that the matrix is shared among all UCEs, and thus inferred from all sites in a set of sequences.

**Figure 2.**
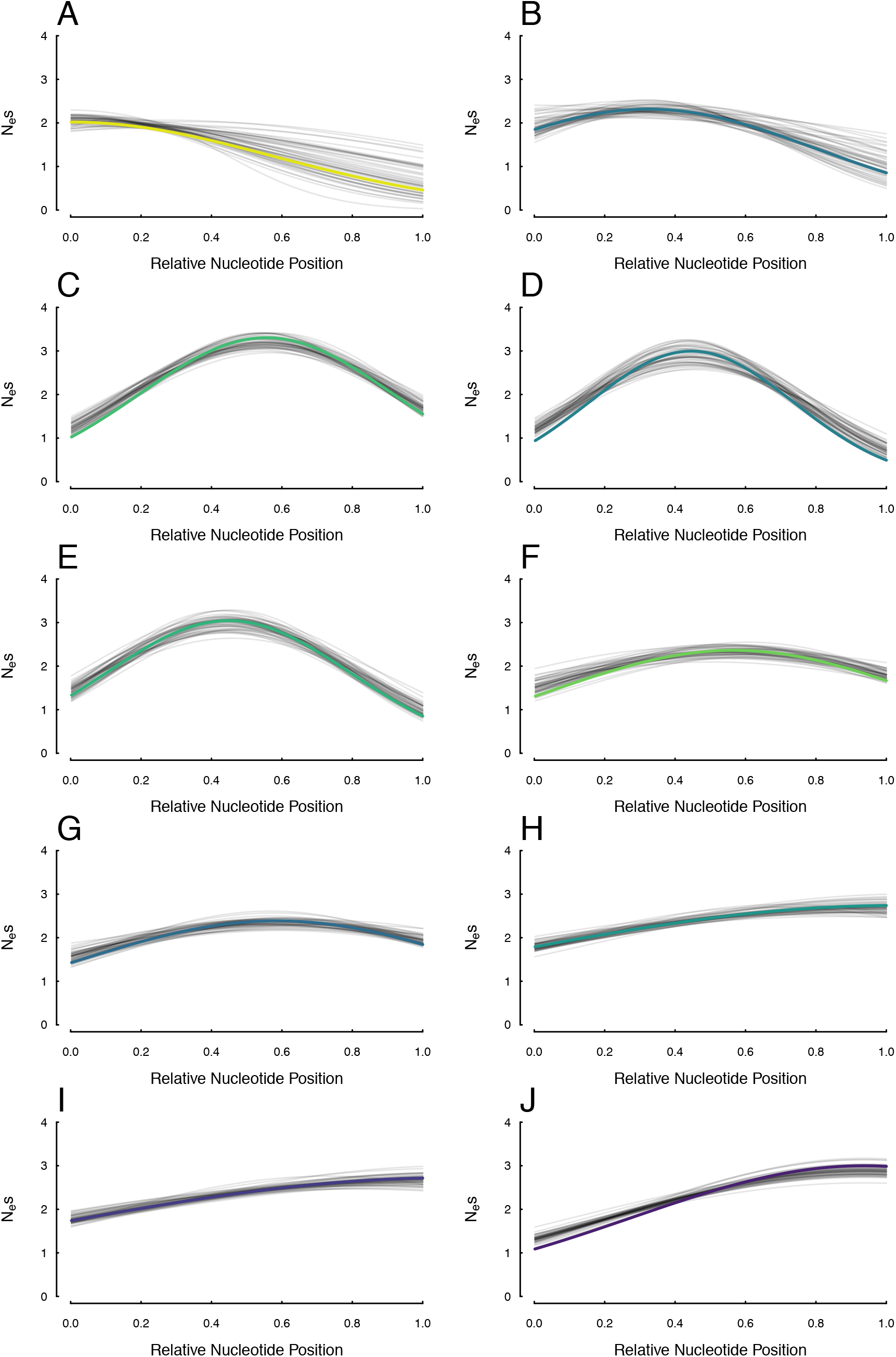
Estimates of the centering, width, and magnitude of the sensitivity to selection distributions for 10 of the 22 simulated UCEs under the SelON model from the simulation shown in Fig. 3A. The thick colored lines represent the generating distribution, with each simulated replicate shown in transparent black lines. The colors are consistent with the color of the points depicted in Fig. S2-3.

In each of the simulation scenarios we examined, the SelON model fit the simulated data substantially better than a standard GTR+*Γ* partitioned across each UCE. The average 4AICc improvement was 5583.9 AICc units across all the simulation replicates and simulation scenarios (the average model weight for SelON = 0.9999), and there was never a single replicate under any scenario in which the GTR+*Γ*model fit better. Aside from improvements in the overall fit, perhaps the most striking difference between the SelON and GTR+*Γ* models were their respective branch length estimates (Fig. 3). It is first worth noting that SelON had difficulties in properly estimating the lengths of the two descendant branches from the root (see Fig. S4). We suspect this is partly an identifiability issue, as arbitrarily rerooting the tree along any portion of one of these branches had a trivial impact on the overall likelihood. Currently, we are concerned that, unlike other non-reversible models (e.g., Huelsenbeck et al. 2002; Zou et al. 2012; Klopfstein et al. 2015), SelON may be incapable of locating the root of a tree and so the inclusion of an outgroup is recommended to absorb the impact of these potential identifiability issues. When following this procedure, SelON correctly identifies the location of long and short branches within the so-called “ingroup”, even when they are distributed among distantly related taxa (e.g., Fig. 3; Fig. S4). However, the overall tree lengths, which were determined by summing the branch lengths in a given tree, were consistently 10% longer than the generating trees across both scenarios. By contrast, the branch lengths inferred by GTR+*Γ* from these same data strongly underestimated the true evolutionary distances among taxa by roughly 50% (Fig. 3C,D; 0.60 in Scenario 1, and 0.49 Scenario 2).

**Figure 3.**
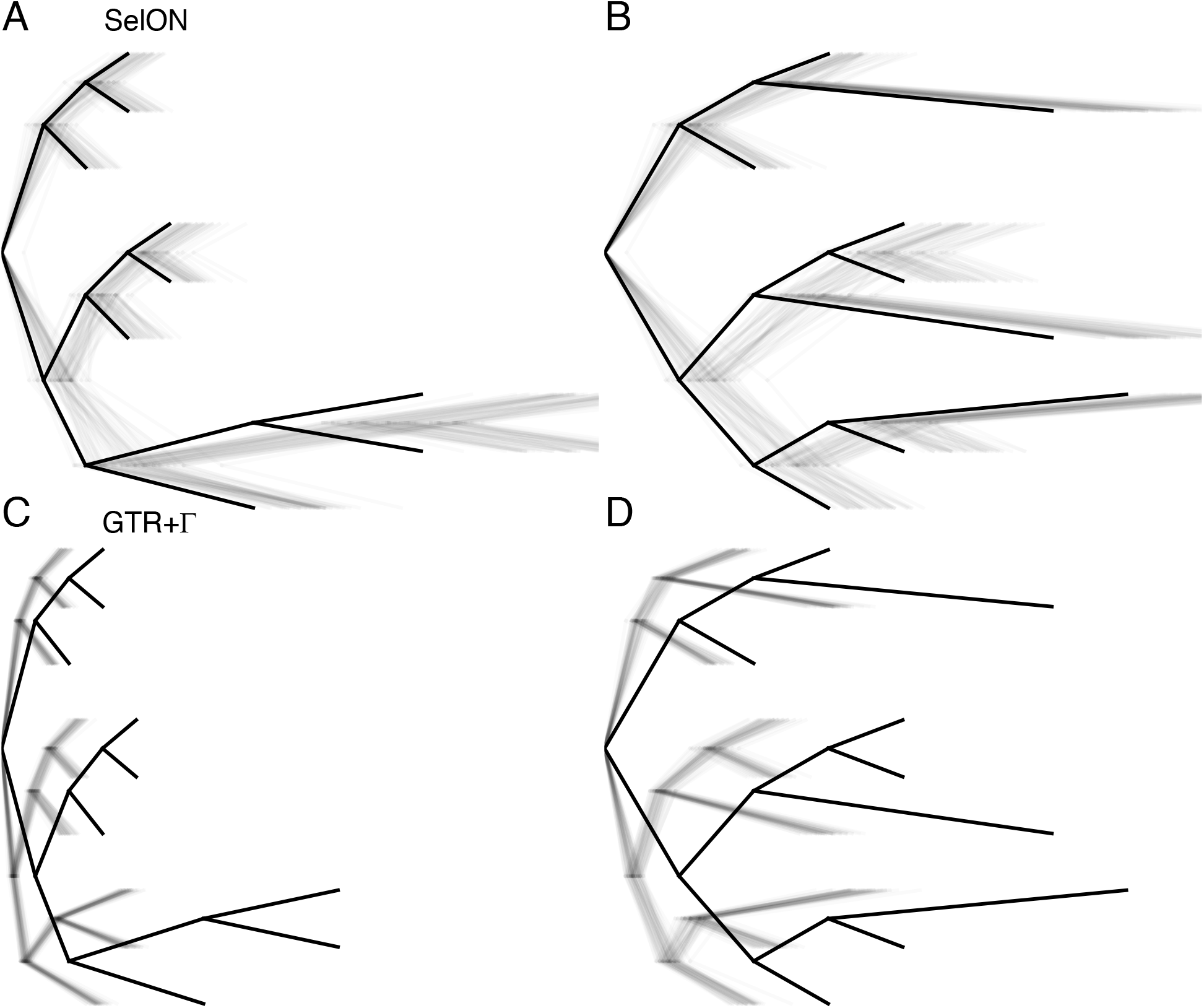
The two scenarios to test whether the distribution of long branches (0.10 expected substitutions/site) and short branches (0.025 expected substitutions/site) within a tree can impact branch length estimates. The first and second row represents the estimated branch lengths inferred under SelON (panels A and B) and GTR+*Γ* (panels C and D), respectively. The tree with the thick branches represents the generating model, and each transparent line represents the estimates from an individual simulation. In all cases, the trees were estimated with a single outgroup taxon that was removed. SelON performed reasonably well, whereas the branch lengths inferred by GTR+*Γ* strongly underestimated evolutionary distances among taxa.

We fit our SelON model to an empirical data set comprising 400 randomly selected nuclear UCE loci used (Crawford et al. 2012) to determine the predominant signal in the placement of turtles relative to archosaurs (bird+crocodiles) and lepidosaurs (lizards+tuataras). These data found overwhelming support for turtles being sister to archosaurs, as opposed to being sister to lepidosaurs (i.e., “Ankylopoda” hypothesis) which previous analyses of limited microRNA data had suggested (see Lyson et al. 2012). It is important to note that in both Crawford et al. 2012 and our study we found the branch lengths differentiating the two models to be extremely short (Figure 4A). When we compared the topologies consistent with these competing hypotheses, by fitting a standard GTR+*Γ*model partitioned across the 400 loci, we also found strong support for the turtle-archosaur alliance (four-*Γ*categories: ln*L_AH_* = – 427,732.0) over the Ankylopoda hypothesis (ΔAICc=180.85, four-*Γ* categories: ln*L_AH_* = −427,822.5). The inferred branch lengths were also strongly consistent with those inferred by Crawford et al. (2012), with the snakes and lizards showing long branches relative to the rest of the tree (Fig. S5). When SelON was fit to these same data, not only does SelON provide an extraordinary improvement in overall fit compared to GTR+*Γ* (*Δ*AICc=85,701.7), but it also indicated stronger support for the Ankylopoda hypothesis (*Δ*AICc=22.5, ln*L_AH_* = −213,629.8, vs. ln*L_TAA_* = −213,641.1; Fig. 4, S5), but with a very short branch (Fig. 4A). The parameters inferred under SelON implied that there was variation in the magnitude of the sensitivity to selection across the 400 UCEs, with the shapes generally following the expected Gaussian distribution with regions most sensitive to selection often being centered near the middle of a given sequence (Fig. 4C).

**Figure 4.**
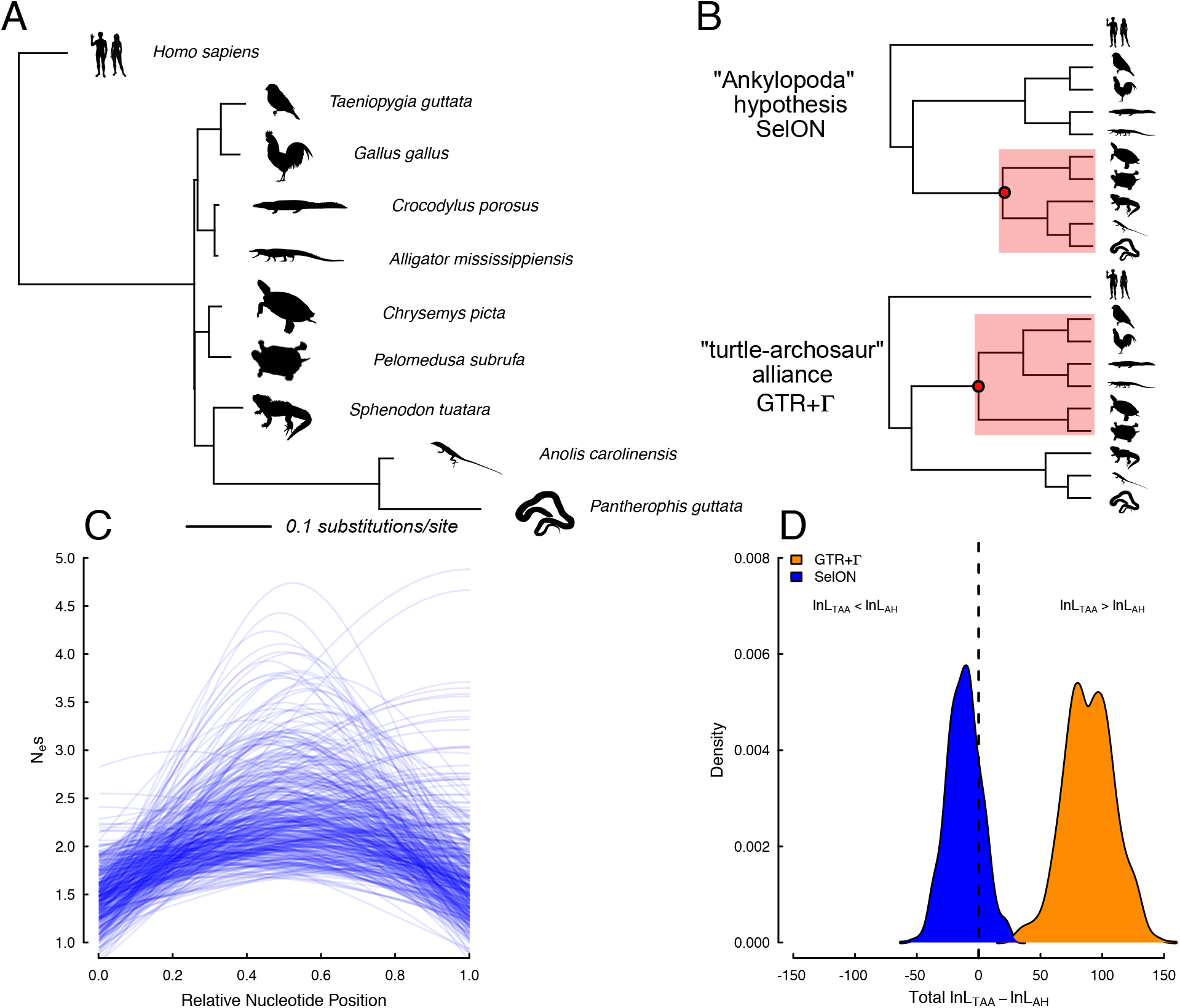
A maximum likelihood phylogram estimated from 400 nuclear UCE loci from Crawford et al. (2012) for determining the placement of turtles relative to archosaurs (bird+crocodiles) and lepidosaurs (lizards+tuataras). (B) When comparing topologies, there was overwhelming support under GTR+*Γ* for turtles being sister to archosaurs (turtle-archosaur alliance), whereas under SelON, not only does it provide an extraordinary improvement in overall fit compared to GTR+*Γ*, but it also indicated stronger support for the Ankylopoda hypothesis (i.e., turtles sister to lepidosaurs). (C) The inferred *N_e_s_max_* shows there is variation in the magnitude of the sensitivity to selection across the 400 UCEs, and their concomitant shapes generally followed the expected Gaussian distribution where regions most sensitive to selection are centered near the middle of a given sequence. (D) The results of a bootstrap procedure, which sampled with replacement the log-likelihood differences between the two competing topologies across the 400 UCEs, found that 81% of the data sets under SelON still maintained support for the Ankylopoda hypothesis, and with GTR+*Γ* 100% of the data sets maintained support for the turtle-archosaur relationship. All silhouettes come from http://phylopic.org.

To better understand the discrepancy between GTR+*Γ* favoring the turtle-archosaur relationship, and our SelON model favoring the Ankylopoda hypothesis with the same data, we examined the loglikelihoods of each individual UCE. The absolute support of one topology over the other under either model was more or less split evenly among our set of UCEs. Of the 400 UCEs used, there were 109 favoring the turtle-archosaur relationship, and 103 favoring the Ankylopoda hypothesis under both models. There were 93 UCEs that favored the turtle-archosaur relationship under GTR+*Γ*, but switched to supporting the Ankylopoda hypothesis when fit under the SelON model, and 95 UCEs with the opposite pattern. Performing a simple bootstrap procedure, where we sampled with replacement entire UCEs and summed the log-likelihood differences between the two competing topologies from the sampled UCEs, we found that 81 % of the pseudoreplicate data sets under SelON maintained support for the ML topology, whereas with GTR+*Γ* 100% of the pseudoreplicate data sets maintained support for the turtle-archosaur relationship (Fig. 4D).

From a model fit perspective, which we measured using AICc, SelON fits better than GTR+*Γ* and so the inferred topology and inferred branch lengths should, in theory, better reflect the true evolutionary distances among these taxa. However, whether the SelON model produces data sets that better resemble the observed data is a separate, but equally important, question (see, for example, Goldman 1993; Bollback 2002; Brown 2014; Beaulieu et al. 2019). We performed a simple test of model adequacy. This involved simulating 400 UCEs with the same sequence lengths as the empirical ones, under both SelON and GTR+*Γ* models, using the MLE parameters estimates on their respective MLE topologies and branch lengths, from which we measured similarity to the observed sequences. Surprisingly, SelON and GTR+*Γ* performed equally well -- that is, on average, the simulated alignment matrix under each model was roughly 85% identical to the observed sequences. Thus, both models adequately described the underlying heterogeneity structure of a given UCE.

It is unsurprising that the *Γ* rate distribution can handle the spatially structured heterogeneity of UCEs (Yang 1994). However, it is important to emphasize that the shape of the discrete *Γ* rate distribution is defined by single parameter, *α*, optimized based on information coming from all sites, with the overall likelihood of an individual site being the average likelihood for that observation under each rate category. In other words, rather than summing the log likelihoods using only the rate category with the best likelihood at each site, the “best” rate is really a weighted average based on the fit of the different rate categories. Thus, it is possible that the rates used for generally invariant sites in the middle of a UCE may be affected by the inclusion of more variable sites at the ends that require higher rate categories to accommodate their higher rates. Interestingly, under GTR+*Γ* there was a significant positive relationship (slope=0.456, *P*=0.045; Fig S6) between the average *Γ* rate for each individual UCE and topological support for turtle-archosaur relationship (see Fig. 5, S7). This was also consistent when examining the site-wise distribution of topological support based on position from the most conserved center, where the cumulative support under GTR+*Γ* for the turtle-archosaur relationship increases as one moves towards the ends. Again, presumably, this is due to the presence of sites with the highest rates, which also showed the strongest support for the turtle-archosaur relationship. We also note that these results are consistent even when comparing topological support based on estimating branch lengths and fitting GTR+ *Γ* to each UCE individually (see Fig. S6, S8). This suggests that the preference for one topology over the other is likely due to differences between the two models.

**Figure 5.**
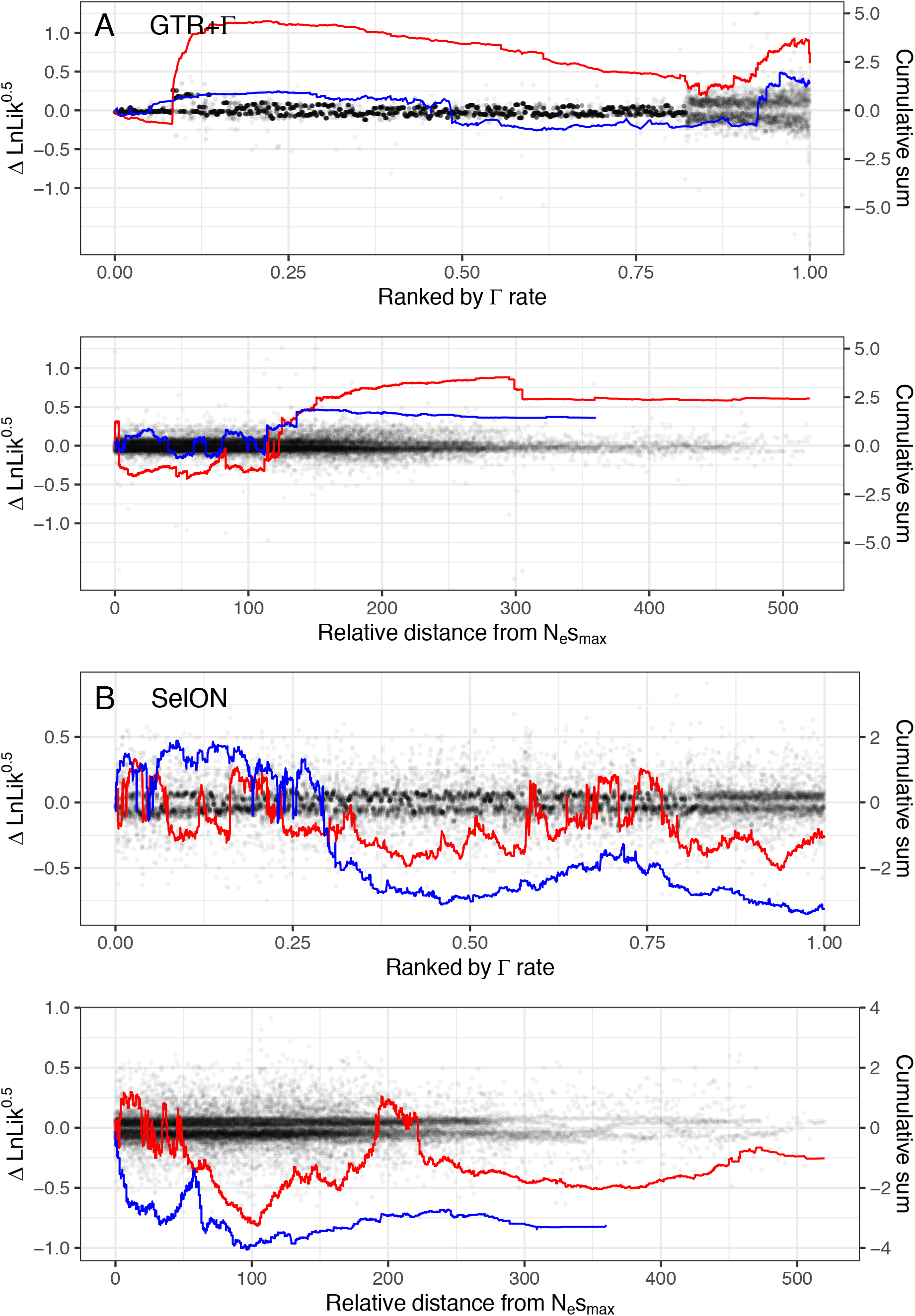
The site-wise patterns of topological support (support defined as Æn*L*=ln*L*_TAA_ – ln*L*_AH_) for 50 randomly selected UCEs, first fit with GTR+*Γ* and SelON (red lines), and then refit, but with 50% of their sites trimmed, targeting the more variable portions at each end (blue lines). With GTR+*Γ*, trimming the variable ends had a marked effect on reducing the support for the turtle-archosaur relationship, both in terms of site distance from the most conserved center (determined by the location of the inferred *N_e_s_max_* under SelON using the untrimmed data) and, most importantly, for a given weighted-average *Γ* rate at a site. With SelON, overall support for the Ankylopoda hypothesis (i.e., lines generally falling below 0) actually increased with the trimmed data set, but overall, tended not to vary substantially and generally followed the same pattern of support as the full data set.

To investigate this pattern even further, we randomly sampled 50 UCEs, first fitting SelON and GTR+*Γ* models to each topology, and then refitting the same models to the same UCEs, but with 50% of their sites trimmed, targeting the more variable portions at each end. (note that the proportion of sites trimmed from a given end depends on the location of the ultraconserved region; see Material and Methods). With GTR+*Γ*, trimming had a marked effect on the support for the turtle-archosaur relationship, both based on site distance from the most conserved center and for a given weighted-average *Γ* rate at a site (Fig. 5). In fact, trimming had the effect of making support for one topology over the other become more equivocal under GTR+*Γ*. With SelON, overall support for the Ankylopoda hypothesis actually slightly increased with the trimmed data set, but overall, tended not to vary substantially and generally followed the same pattern of support as the full data set (Fig. 5). Given that likelihood models can be inconsistent with wrong rates (Kolaczkowski and Thornton 2004), the fact that the width of the flanking region can affect rates throughout a UCE under a GTR+*Γ*, but not SelON, suggests the former may be more prone to issues such as long branch attraction.

## Discussion

Ultraconserved elements (UCEs) are important loci for phylogenetic reconstruction. UCEs consist of a highly-conserved core that makes them easy to align across a divergent set of taxa (such as humans and birds), and their increasingly variable flanking regions are assumed to contain phylogenetic information across a variety of evolutionary timescales. From a mechanistic standpoint, the conserved nature of UCEs implies that these regions are under strong selection that, in turn, maintains both deep and shallow homologies (Bejerano et al. 2004; Woolfe et al. 2005; Katzman et al. 2007). Our work is the first step in analyzing UCEs using explicit and spatially varying models of stabilizing selection.

Currently, the most widely used substitution-based nucleotide models, such as the GTR model with a discrete *Γ* rate distribution (+*Γ*), simply describe the relative rates of change between nucleotide bases. In the context of UCE evolution, the GTR+*Γ* model is not based on any explicit model of natural selection and, as a result, ignores the spatial structure of UCEs. This forces the model to impose an inappropriate rate distribution, resulting in the presence or absence of high rate sites affecting the support for particular topologies at more conserved sites (Fig. 5, S6-7). One approach to mitigating this effect would be through a more sophisticated partitioning of the rate variation across the set of UCEs (Tagliacollo and Lanfear 2018). Another approach is to explicitly model the spatial variation in natural selection along a UCE. Taking the latter approach here, we model the nucleotide substitution process of a UCE, which includes the effects selection, mutation, and drift processes. In addition to being a better fitting model than the standard GTR+ *Γ*, the spatial structuring in the variation we impose in the substitution process is generally robust to the sequence length and degree of variation contained within a single UCE. Further, because selection is modeled explicitly, testing more sophisticated, model-based hypotheses of how selection varies spatially within and between UCEs is relatively straightforward.

Phylogenetic inference with an inadequate model can lead to the incorrect tree inference (Felsenstein 1979; Kolaczkowski and Thornton 2004), and so, the discovery that model choice affects phylogenetic position of lineages (e.g., the placement of turtles relative to archosaurs and lepidosaurs) suggests that this has real world consequences. Perhaps more fundamentally, our results emphasize the potential issues with the continual pursuit of candidate loci that capture multiple levels of relationships (e.g., Shedlock and Okada 2000; Edwards et al. 2017; Dornburg et al. 2019). As we show, that in combination with an inadequate model, the rate variation of a UCE does not structure cleanly across timescales. In other words, under GTR+*Γ* the increasingly variable flanking regions do not speak exclusively to the shallow portions of the tree and have a clear effect on the deeper splits as well, and so investigator choice of where to cut off a UCE may have a substantial effect if they use a GTR+*Γ* model. We recognize that our analyses and interpretations are somewhat biased towards defining what is “ultraconserved” based on parameter estimates from our new and relatively untested SelON model. Nevertheless, our results highlight a need to better understand, and better account for, the conflicting signal in these flanking regions, which exhibit generally higher substitution rates that have a disproportionate impact on the signal (i.e., whether turtles are sister to archosaurs or lepidosaurs). Our concerns over the impact of highly variable regions are not completely limited to UCEs. They also extend to historically used markers fundamental to systematic pursuits, such as the small ribosomal subunits (e.g, SSUs like 18S rRNA, or other parts of the ribosomal operon) and new high-throughput approaches that target conserved regions, but also capture linked and/or flanking non-target sequences (e.g, intronic sequences for exon capture; see Johnson et al. 2019).

Our finding that turtles are placed as sister to lepidosaurs was based on a concatenation approach, where the overall likelihood of topology comes from a simple summation of the log-likelihoods across all the UCEs in the set. We fully acknowledge that this may not be ideal, as gene trees do not necessarily always match species trees (Maddison 1997). Indeed, the incredibly short branches that differentate the two hypotheses suggest the divergence of these lineages occurred very rapidly which, in turn, would likely lead to a large amount of incomplete lineage sorting. Nevertheless, we did look at the signal for each topology for a given UCE by examining the absolute support for one topology over the other. This is, of course, different from conducting a full gene tree-species tree analysis. Nevertheless, whether a concatenation approach or a gene tree approach is used, having an appropriate model that best fits the evolution of sites within loci of interest is important.

The UCE specific parameters estimated by SelON provide a compact way of quantifying the strength and variation in selection within and across UCEs. By contrast, classic models of nucleotide substitution are hard to interpret biologically, at least in the context of UCE evolution. When the rates of substitution simply reflect mutation, then models within the GTR family are consistent with models of neutral evolution. Even if GTR seems to behave phenomenologically like an explicit model of consistent, stabilizing selection there is no inherent bias towards any particular nucleotide at a given site at any given moment. However, the utility of the +*Γ* extension is that it allows for evolution at rates above and below those expected under neutrality. While GTR+*Γ*adequately generates data sets very similar to the observed data sets, the long-term expectations under the model is that eventually the highly-conserved nature of the UCE will break down. In other words, the model views UCEs as containing regions with very low rates of neutral evolution that have not yet reached equilibrium.

Thus, in our view, in order to invoke stabilizing selection as the evolutionary force underlying the GTR+*Γ*model one must assume infinite, but shifting stabilizing selection rather than finite, but varying stabilizing selection as the way the model is usually interpreted. That is, under GTR+*Γ*, variation in substitution rates between sites does not reflect variation in the strength of stabilizing selection. Instead, the variation in substitution rates represents variation in the rate at which the optimal nucleotide shifts. Furthermore, in order for the sequence to track this change on time scales consistent with the model, the strength of this stabilizing selection must be consistently strong across all sites such that the substitution occurs almost immediately after the shift in the optimal nucleotide (see Beaulieu et al. 2019 for a similar argument about models of codon evolution). This interpretation is analogous to fitting models of trait evolution where even though evolutionary change for a focal trait is consistent with Brownian motion, it is also correlated with another factor and/or trait that is following a continuously shifting optimum (see Beaulieu et al. 2012).

While SelON extends models by allowing the strength of selection to vary spatially, in its current form, it assumes that sequences start out already in equilibrium and that the optimal nucleotide sequence does not change along the tree. This means that, unlike GTR+*Γ*, the general UCE structure is expected to persist indefinitely. Future extensions of SelON could easily allow the optimal sequence to vary among lineages and over evolutionary time through the use of a hidden Markov modelling approach (e.g., Galtier 2001; Beaulieu et al. 2013; Beaulieu and O’Meara 2016). Alternatively, one could relax our current assumption that the magnitude of stabilizing selection for the optimal nucleotide is solely determined by its position. In this case, one could treat the position dependent strength of selection at a site as an expectation of a random variable from some distribution, rather than as an exact value as we do here. For example, the strength of stabilizing selection for a given site could be modeled as if it were drawn from a Gamma distribution whose expected value varies spatially according to a Gaussian function (i.e. SelON+*Γ*). In fact, a combination of these model extensions under the SelON framework should be able produce a range of models with behavior similar to the standard GTR+*Γ* but that is actually more consistent with how this model is commonly perceived. Generalizing our model in this manner would allow for testing specific hypotheses about the fundamental and poorly understood biological factors contributing to the evolution of UCEs. In addition, these model extensions should allow us to more effectively extract phylogenetic information on shallower branches by including additional flanking regions beyond the core of a given UCE that are currently ignored.

Despite intense study of UCEs, we still know very little about the mechanisms responsible for their persistence across deep evolutionary time and the factors contributing to their rate heterogeneity. Given the fact that the spatial variation in nucleotide conservation is a common property of UCEs, it should not be surprising that a Gaussian model of stabilizing selection, which employs three parameters to describe the center of the UCE, the strength of selection at the center, and how quickly selection decreases away from the center, fits better than the GTR+ *Γ*; a single parameter random effects model that implicitly, rather than explicitly, models the effects of selection. Looking further ahead, one could imagine testing a range of spatial functions, such as a truncated or bimodal function, rather than exclusively assuming the strength of stabilizing selection follows a Gaussian function. Fitting other spatial functions to UCEs should allow finer grade differentiation of UCEs. Better differentiation of UCEs should both improve phylogenetic reconstruction of evolutionary history and help researchers exclude or even identify the sources of selection underlying UCE evolution.

## Material and Methods

### Implementation

We implemented our SelON model, as well as carried out all subsequent simulations and empirical analyses, in our R package *selac* (Beaulieu et al. 2019). As input, all that is required is a directory containing fasta files of the individual UCE sequences for a set of taxa, and a phylogeny depicting the hypothesized relationships among them (due to computational constraints only fixed topologies are optimized). The starting values for 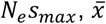, and *σ* that define the continuous shape function are chosen at random, and we start with a global mutation rate matrix that assumes all rates are 1 (i.e., we start with a Jukes-Cantor substitution process). The initial optimal nucleotides, 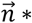, are based on the majority rule, where the most observed nucleotide at each site is considered the starting optimum. Since the branch lengths are shared among all sequences, the starting values for the branch lengths come from the estimates under the F81 model (Felsenstein 1981) of substitution using the R package *phangorn* (Schliep 2011).

To optimize branch lengths and model parameters, we employ a four-stage hill climbing algorithm. The first stage optimizes the branch lengths, but because our model is non-reversible, we are unable to make use of the standard fast re-rooting algorithm that is commonly implemented for estimating branch lengths (Felsenstein 1981). Instead, we devised an alternative procedure that first optimizes the particular order of branch length “generations”, starting with the terminal branches being optimized, followed by the branches that subtend them, and so on, until we reach the root. The order here is important, because in effect we are always optimizing the length of the branches that subtend increasingly inclusive clades, whose descendant branch lengths have already been optimized in previous generations. Once we have optimized all branches in the tree, this does not guarantee that each branch is at their optimal length. So, we repeat this cycle until either nine additional cycles have been conducted, or there is a <1% difference in the log-likelihoods between successive cycles.

The second stage of our algorithm optimizes the UCE-specific parameters 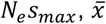, and *σ* that set the shape of the continuous sensitivity to selection function, and the third stage optimizes the mutation rate parameters, *μ_ij_*, shared by all UCEs in the set. In each of these stages, parameter optimization is carried out using a bounded subplex routine implemented in the NLopt library, and made available for R through the package *nloptr* (Johnson 2019). The fourth stage optimizes the optimal nucleotide sequence, 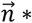, for each site in each UCE in the set. Our four-stage cycle is repeated until either 11 additional cycles have been conducted, or there is a <1% difference in the overall log-likelihoods between successive cycles. This process is slow, and it is likely that others could develop faster optimizations if they incorporate this model into other software.

### Simulations

We evaluated the performance of SelON by simulating UCEs using two different nine- and ten-taxon topologies under SelON (Fig. 3; Fig. S4), and estimating the bias of the inferred model parameters and branch lengths from these data. First, the ten-taxon topologies are the nine-taxon topology with a single outgroup taxon added to test the impact of whether adding an outgroup improves branch lengths estimates for the nine-taxon ingroup (see *Results* in the main text). Second, the different topologies reflect ways in which long branches (0.10 expected substitutions/site) and short branches (0.025 expected substitutions/site) are distributed within a tree. The purpose was to determine not just the behavior of our SelON model, but also the behavior of the branch length estimates from standard models of nucleotide evolution (i.e., GTR+*Γ*) when the true underlying model assumed varying degrees of stabilizing selection. The generating model was based on the parameter estimates and site lengths from 22 randomly selected UCEs from our full analysis of the turtle dataset (see *Placement of Turtles)* fit under SelON, which, cumulatively, produced data sets with 10,000 sites. We conducted 50 replicates of each simulation set, pulling ancestral states from the equilibrium base frequencies for each site. To ensure robustness of the parameter estimates, we optimized using the generating model parameters as starting points, as well as repeated the model optimization three times with different naive starting points with respect to the UCE-specific parameters. When fitting the GTR+*Γ*model of nucleotide evolution the *Γ*-distribution was approximated using the generalized Laguerre quadrature method with four categories (Felsenstein 2001).

### Placement of turtles

We fit our SelON model to a published data set of 1,145 UCEs and their variable flanking DNA (Crawford et al. 2012). These loci were used to determine whether the phylogenetic position of turtles is sister to archosaurs (birds+crocodiles), or sister to the lepidosaurs (lizards+tuatara; “Ankylopoda hypothesis”, see Lyson et al. 2012), which we reevaluated here. However, due to computational limitations, rather than using all 1145 UCEs we randomly selected 400 from this broader set. To ensure robustness of the parameter estimates, we repeated the model optimization three times, each with different naive starting points with respect to the UCE-specific parameters. In addition to our SelON model, we also fit two GTR+*Γ* models partitioned by UCE (as was done in the original study) where the *Γ* distribution was approximated by the generalized Laguerre quadrature (Felsenstein 1981). Two separate analyses were conducted, one where the number of rate categories was set to four categories, and another where the number of rate categories was set to eight (in no case did adding extra categories improve the fit from the standard four category *Γ*-distribution). For all model comparisons between SelON and GTR+*Γ* fits we rely on sample size corrected AICc, where the sample size, *n*, is equal to the number of taxa multiplied by the number of sites (see Beaulieu et al. 2019).

In addition to examining improvement in model fit, we also evaluated and compared the model adequacy of SelON and GTR+ *Γ*. This involved simulating 400 UCEs with the same sequence lengths as the empirical ones, under both SelON and GTR+*Γ* models, using their respective MLE parameters, topologies, and branch lengths. We conducted 100 replicates of each simulation set, pulling the starting states from the equilibrium base frequencies for each site. For GTR+*Γ*, the equilibrium base frequencies and *Γ*-rate multiplier for a given site was based on a model-average of the site likelihood across each discrete *Γ*-rate category. Similarity was calculated as the number of matches at any given site for any given taxa divided by the total number sites.

To determine whether our randomly selected sampling from the broader UCE set impacted the support for one topology over another, we also conducted a complementary set of analyses using 50 randomly selected UCEs. We also used this set of UCEs to test the robustness of the support for each topology when trimming a portion of the sites from their variable flanking regions. To do this, we first determined the final size of the trimmed UCE if we removed 50% of the sites, and then determined the stretch of sites of this same size that cumulatively resulted in the lowest overall parsimony score. This procedure ensured that for any given individual UCE, the most conserved portion was retained and that the trimmed sites came from the most variable regions. In other words, a given end may have more or fewer sites removed than another. It all depends on the location of the utltraconserved region. We then refit GTR+*Γ* and SelON to the trimmed data set and compared the resulting overall support for the two competing topologies. We also examined the pattern of site-wise support based on each site’s distance from their respective inferred *N_e_s_max_* (based on the untrimmed UCE analysis) as well as the model-average *Γ* rate.

## Acknowledgements

We thank Andrew Alverson, Brant Faircloth, Jeremy Brown, Cedric Landerer, Alex Pyron, David Swofford, and Russ Zaretski for general discussions and helpful suggestions during various stages of the project. Support was provided in part by NSF award DEB-1355033 (to B.C.O. and M.A.G.) and from the Arkansas Bioscience Institute (to J.M.B.).

