## Supplemental Fig 1-8 for "A spatially-explicit model of stabilizing selection for improving phylogenetic inference"

**This PDF file includes:**

Figures S1 to S8

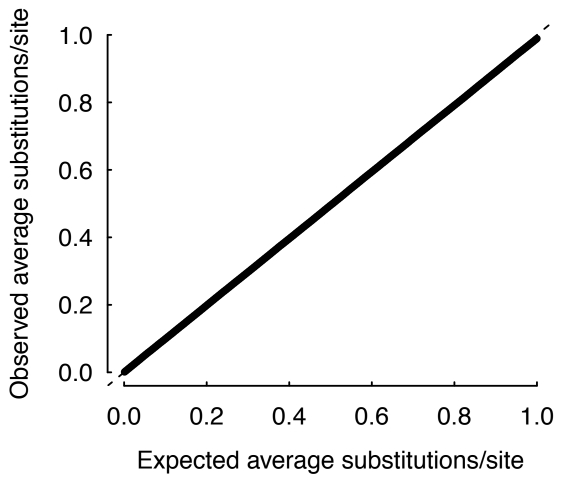


**Fig. S1.** Simulation results demonstrating that one unit of branch length under our SelON model represents one expected substitution per site. The generating model was based on the parameter estimates and site lengths from 22 randomly selected UCEs from our full analysis of the turtle dataset, which, cumulatively, produced data sets with 10,000 sites. We assumed a single starting lineage pulling ancestral states from the equilibrium base frequencies for each site. We then incremented the time by 0.001 expected substitution units and recorded the cumulative number of substitution across sites. We repeated this process 20 times. As would be expected, the expected and observed average substitution per site followed each other exactly.

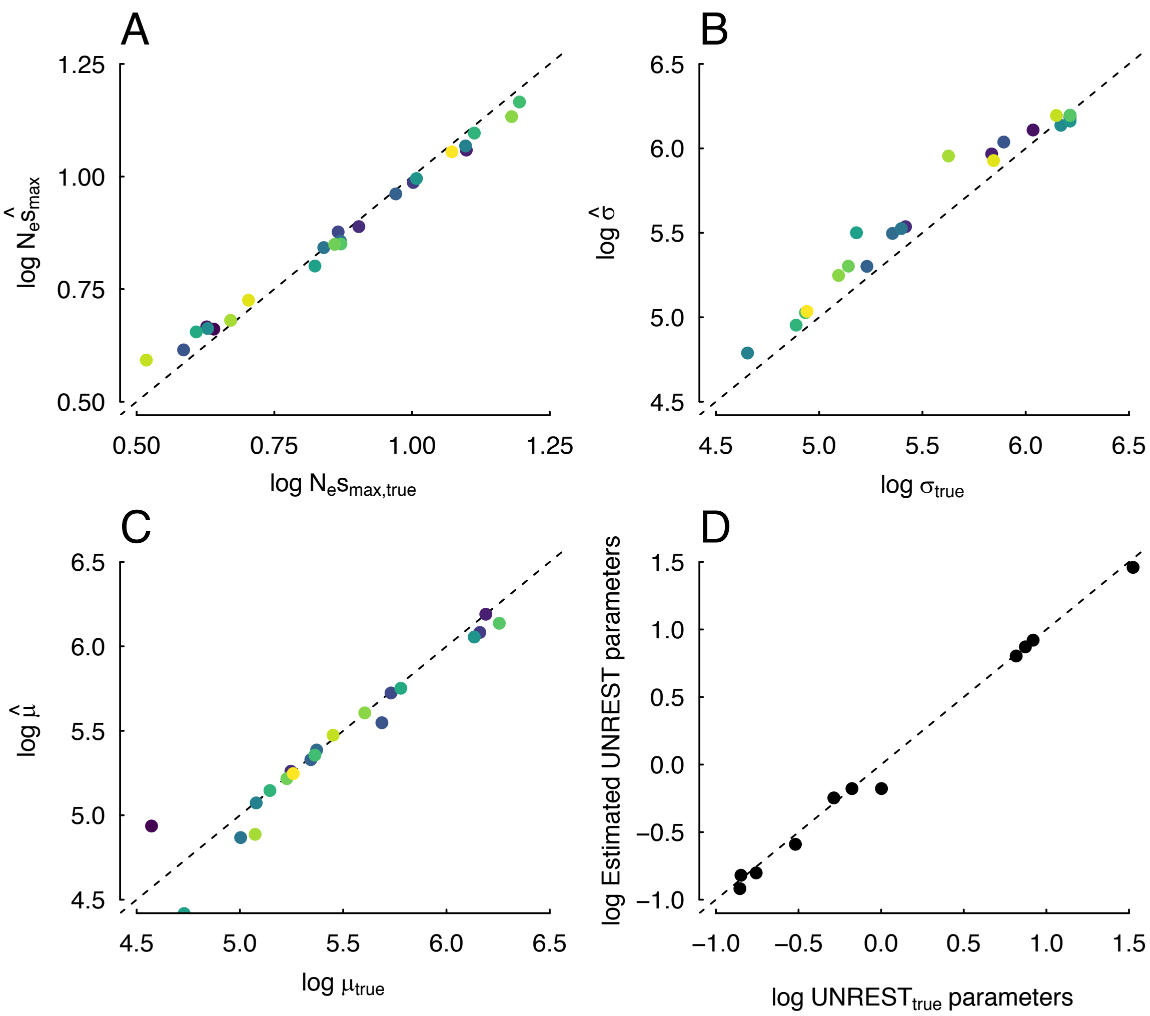


**Fig. S2.** Summary of the simulation where the generating model was based on the parameter estimates from a randomly selected set of 22 UCEs. (A-C) The parameter estimates associated the magnitude, width, and centering of the sensitivity to selection distributions, and (D) the global mutation rates following the UNREST model of nucleotide substitution. The topology used for the simulation is shown in Figure 3A in the main text. The dashed line reflects the 1:1 line.

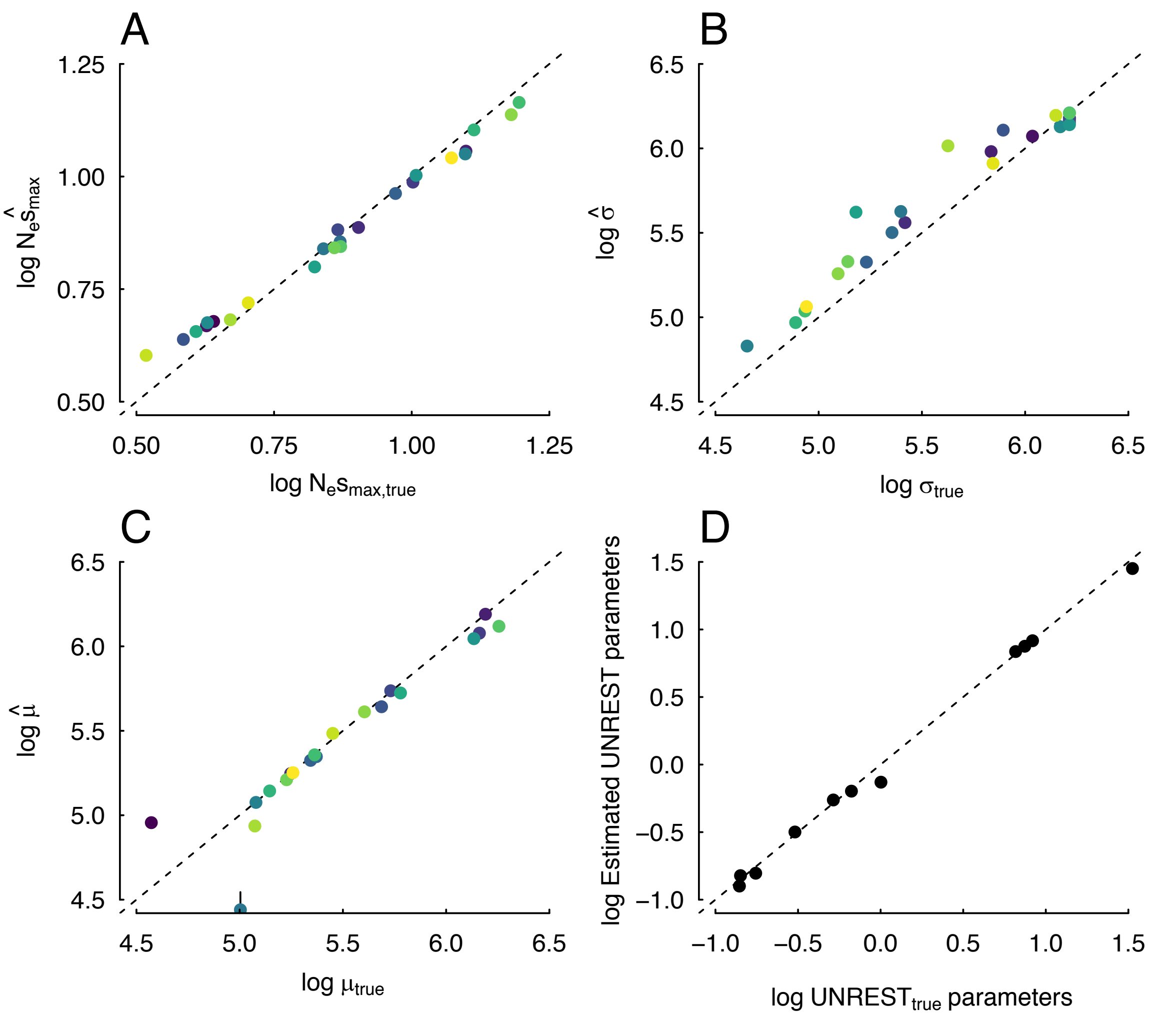


**Fig. S3.** Summary of the simulation where the generating model was based again on the parameter estimates from a randomly selected set of 22 UCEs, but with the topology used for the simulation is shown in Figure 3B from the main text. The dashed line reflects the 1:1 line.

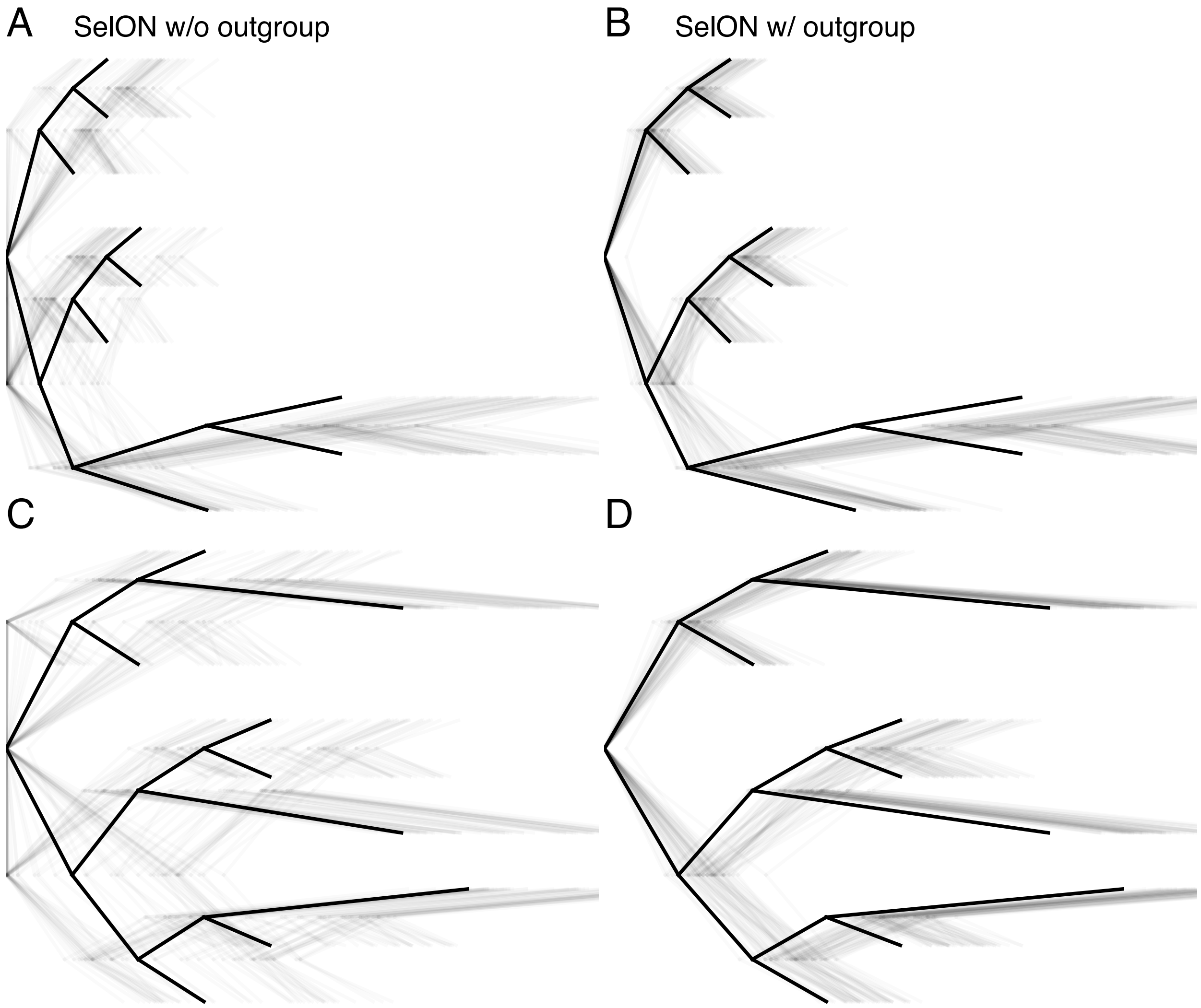


**Fig. S4.** The impact of using an outgroup taxon for two scenarios testing whether the distribution of long branches (0.10 expected substitutions/site) and short branches (0.025 expected substitutions/site) within a tree can impact branch length estimates. The first column represents the estimated branch lengths inferred under SelON without an outgroup (panels A and C) and with SelON with an outgroup that was subsequently removed (panels B and D). The tree with the thick branches represents the generating model, and each transparent line represents the estimates from an individual simulation. We suspect the difficulties SelON had in properly estimating the lengths of the two descendant branches from the root is partly an identifiability issue.

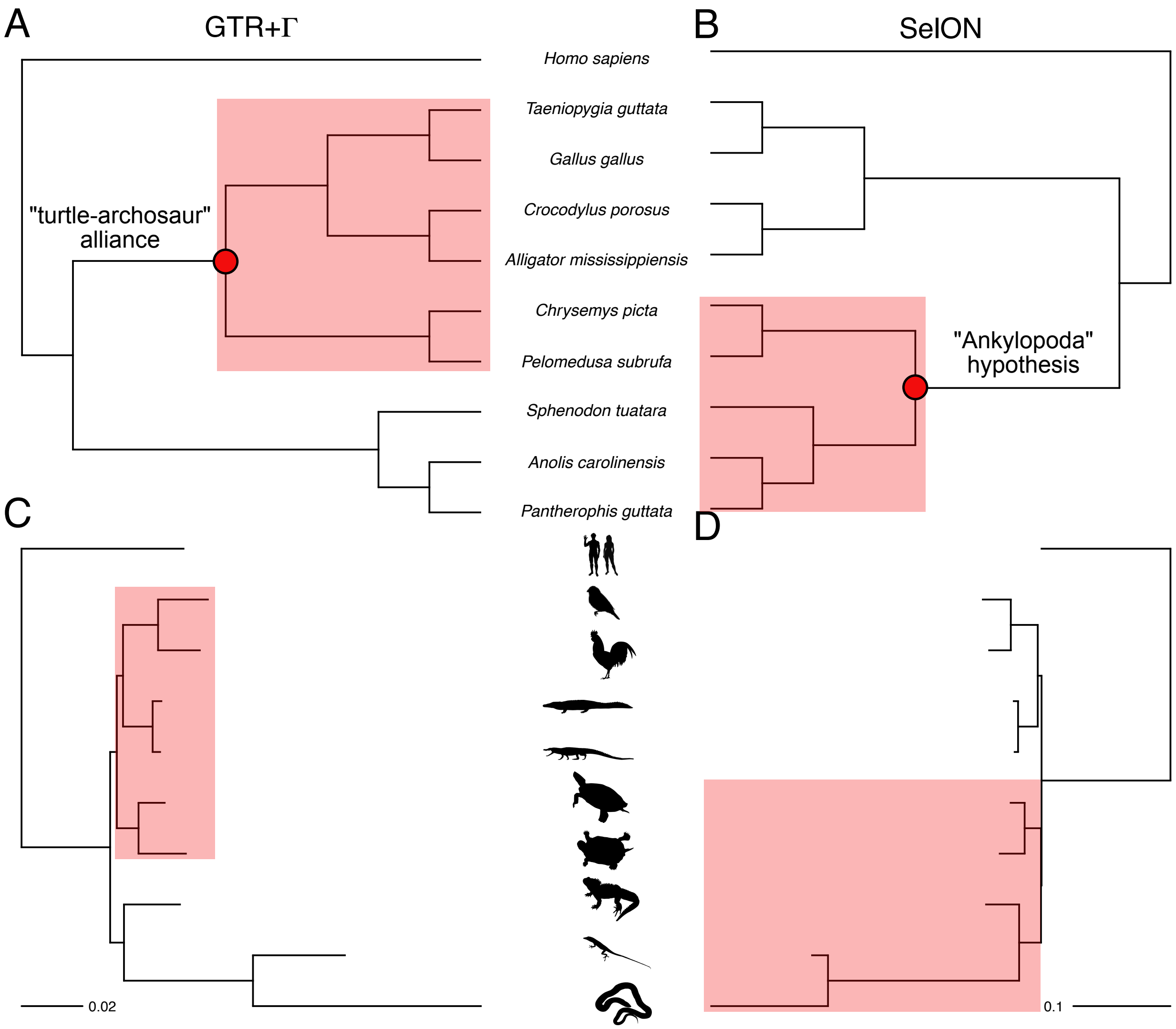


**Fig. S5.** Cladograms and phylograms summarizing the inferred phylogeny for determining the placement of turtles relative to archosaurs (bird+crocodiles, or the "turtle-archosaur" alliance) and lepidosaurs (lizards+tuataras, or "Ankylopoda" hypothesis) under (A,C) GTR+$\Gamma$ and (B,D) SelON. When comparing topologies, there was overwhelming support under GTR+$\Gamma$ for turtles being sister to archosaurs (turtle-archosaur alliance), whereas under SelON, not only does it provide an extraordinary improvement in overall fit compared to GTR+$\Gamma$, but it also indicated stronger support for the Ankylopoda hypothesis (i.e., turtles sister to lepidosaurs).

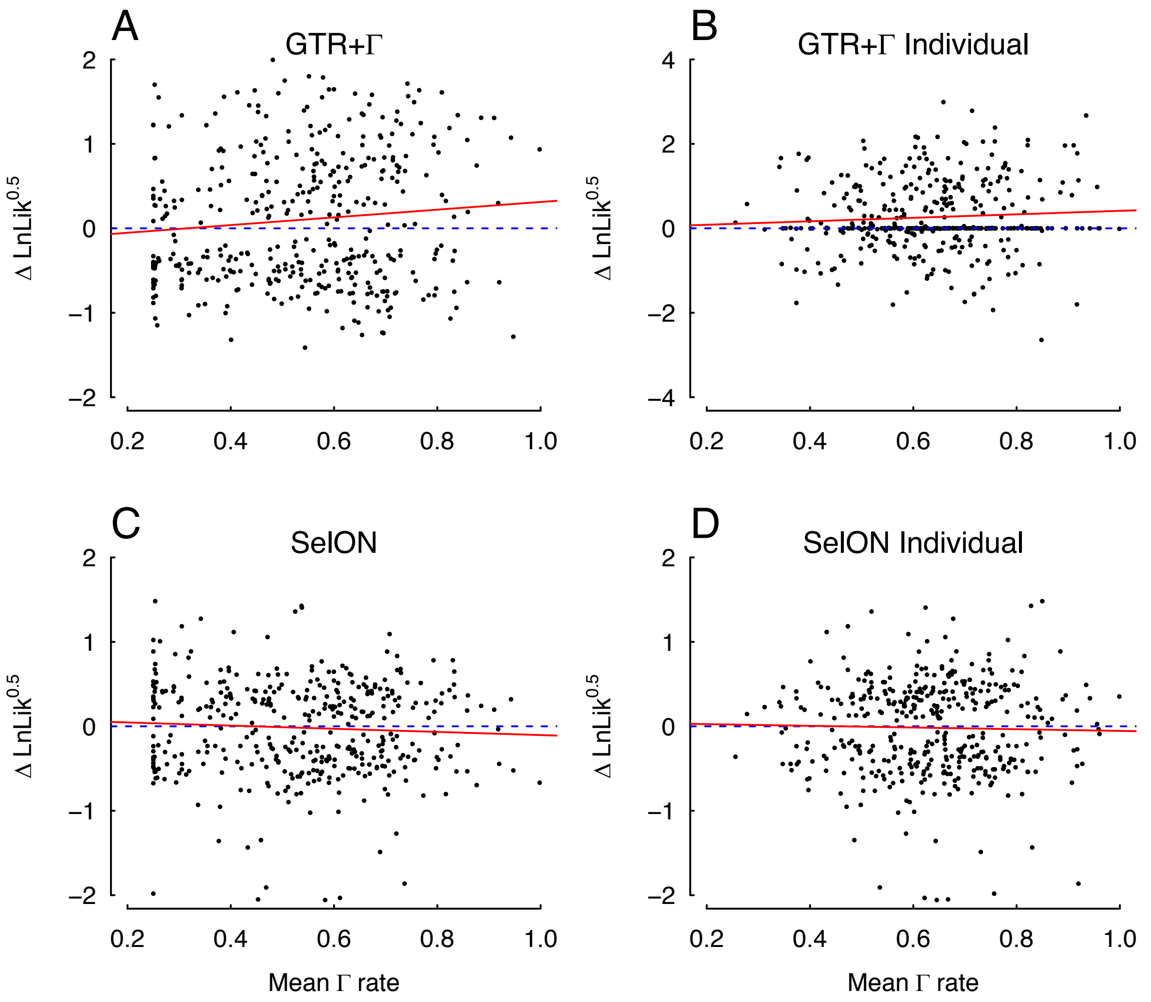


**Fig. S6.** (A,B) A linear regression (red) showing a significant positive relationship under GTR+$\Gamma$ there was (slope=0.456, *P*=0.045; Fig S6) between the average $\Gamma$ rate for each individual UCE and topological support for turtle-archosaur relationship (support defined as $\ln L = \ln L_{TAA} - \ln L_{AH}$). These results are consistent even when comparing topological support based on estimating branch lengths and fitting GTR+ $\Gamma$ to each UCE individually. (C,D) When inferred under the SelON model, the relationship between average $\Gamma$ rate for each individual UCE and topological support for turtle-archosaur relationship is no longer significant. The blue dashed line is a horizontal line at zero.

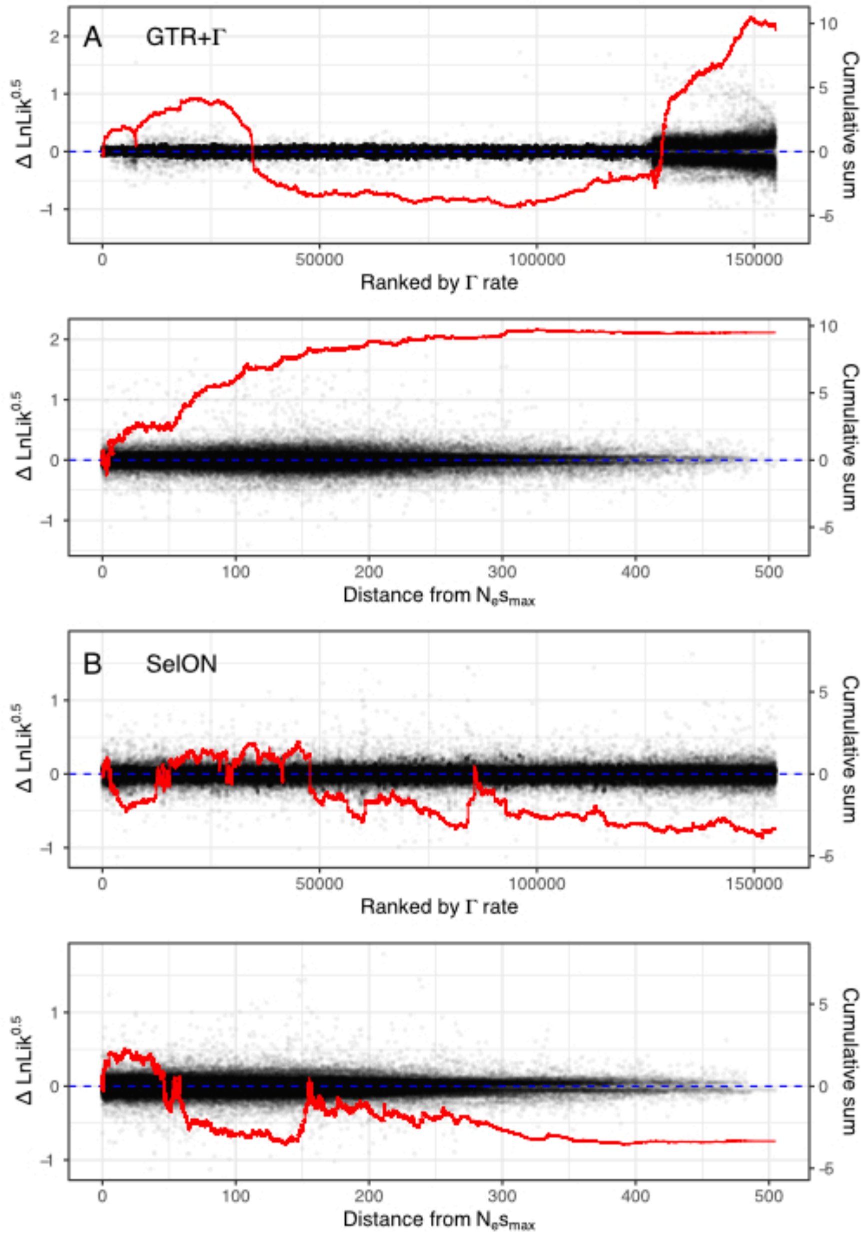


**Fig. S7.** The site-wise patterns (red lines) of topological support (support defined as $\Delta \ln L = \ln L_{TAA} - \ln L_{AH}$) under (A) GTR+$\Gamma$ and (B) SelON model fits to the full 400 UCE empirical dataset. With GTR+$\Gamma$, support for the turtle-archosaur relationship was driven both by the lowest rate sites and highest rate sites, based on a ranking of the weighted-average $\Gamma$ rate at a site. The support also steadily increased as the distance from the presumptive conserved center (determined by the location of the inferred $N_e s_{max}$ under SelON). With SelON, overall support for the Ankylopoda hypothesis was generally supported regardless of weighted-average $\Gamma$ rate and distance from the center.

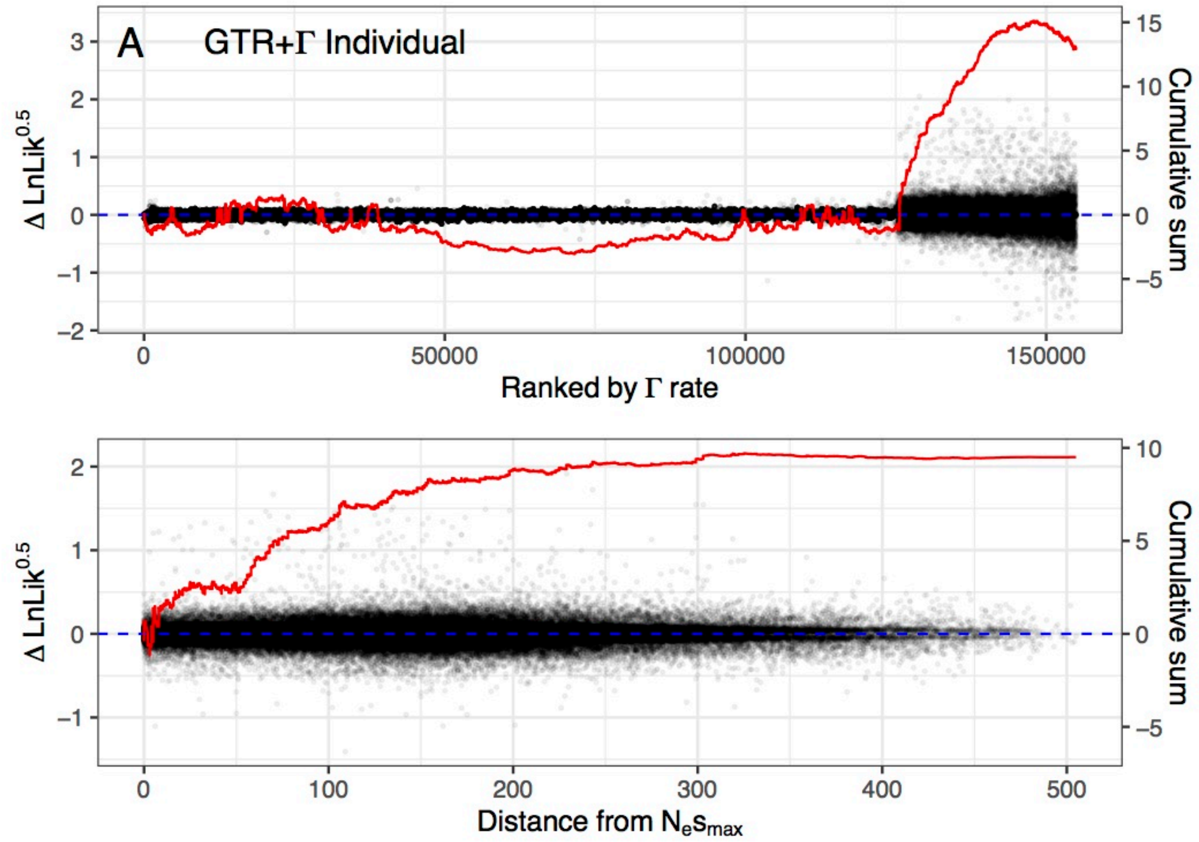


**Fig. S8.** The same site-wise patterns (red lines) of topological support (support defined as $\Delta \ln L = \ln L_{TAA} - \ln L_{AH}$) under GTR+$\Gamma$ model fit as **Fig. S7**, but with branch lengths and model parameters are estimated individually to the full 400 UCE empirical dataset.
